# Cell Type-Agnostic Transcriptomic Signatures Enable Uniform Comparisons of Neurodevelopment

**DOI:** 10.1101/2025.02.24.639936

**Authors:** Sridevi Venkatesan, Jonathan M. Werner, Yun Li, Jesse Gillis

## Abstract

Single-cell transcriptomics has revolutionized our understanding of neurodevelopmental cell identities, yet, predicting a cell type’s developmental state from its transcriptome remains a challenge. We perform a meta-analysis of developing human brain datasets comprising over 2.8 million cells, identifying both tissue-level and cell-autonomous predictors of developmental age. While tissue composition predicts age within individual studies, it fails to generalize, whereas specific cell type proportions reliably track developmental time across datasets. Training regularized regression models to infer cell-autonomous maturation, we find that a cell type-agnostic model achieves the highest accuracy (error = 2.6 weeks), robustly capturing developmental dynamics across diverse cell types and datasets. This model generalizes to human neural organoids, accurately predicting normal developmental trajectories (R = 0.91) and disease-induced shifts *in vitro*. Furthermore, it extends to the developing mouse brain, revealing an accelerated developmental tempo relative to humans. Our work provides a unified framework for comparing neurodevelopment across contexts, model systems, and species.

## Introduction

Single-cell transcriptomics has transformed the study of neurodevelopment, enabling precise mapping of cellular diversity and lineage relationships ^1–5^. Standard approaches identify distinct cell types across datasets ^6–8^, but tracking the maturational state of cells remains a major challenge. While developmental stage can be inferred within individual datasets, generalizable methods for cross-study comparisons are lacking. Establishing robust cell-autonomous and global signatures of neurodevelopment is critical for comparing developmental processes across datasets, experimental models, and species ^9^. Prior efforts have identified low-dimensional representations of developmental stage ^10–12^, but their reproducibility across datasets is unclear. The scarcity of primary tissue samples, coupled with variability in time points and cell types sampled, has hindered meta-analytic comparisons. However, the growing availability of large single-cell datasets from the developing human brain ^13–16^ now offers an unprecedented opportunity to assess the conservation of neurodevelopmental signatures across diverse contexts.

Three primary approaches have been used to infer neurodevelopmental stage: principal component analysis (PCA), differentially expressed genes (DEGs), and pseudo-temporal ordering. PCA applied to bulk RNA-seq often identifies age-associated principal components (PCs) that serve as proxies for developmental stage within a study, enabling comparisons across brain regions, species, and human neural organoids ^11,12,17–19^. DEG-based methods, such as transition-mapping, assess developmental progression by sequentially comparing DEGs between samples of different ages, often using BrainSpan fetal bulk RNA-seq as a reference ^10,20–23^.

Pseudo-temporal ordering reconstructs developmental pathways by arranging cells along latent trajectories ^24,25^, supported by integration, label transfer ^4,26,27^, and dynamic time warping algorithms ^28,29^. While effective within individual studies, these methods lack systematic evaluation across studies and model systems, despite known technical variability in biological and single-cell data. To address this, we leverage 13 developing human brain single-cell RNA-seq datasets ^4,13–16,30–37^, to systematically identify robust tissue-level and cell-autonomous predictors of neurodevelopmental stage.

We find that while tissue composition predicts developmental age within individual studies, it lacks generalizability across datasets. Instead, conserved changes in specific cell type proportions, such as astrocytes and progenitors, provide a reliable cross-study predictor of tissue developmental age. To track cell-autonomous maturation, we develop a cell type-agnostic model that samples genes from diverse developmental processes, enabling robust age prediction across cell types and datasets. This model generalizes to human neural organoids from multiple protocols ^20,38–40^, and embryonic mouse brain development, providing a unified framework for cross-dataset, cross-model, and cross-species comparisons of neurodevelopment.

## Results

We perform a meta-analysis of 13 recent single-cell RNA-seq datasets from the developing human brain, spanning 151 donors across the first to third trimesters, and comprising 2,888,635 cells (**Fig 1**, detailed dataset information in Supplementary table S2). We evaluate the relative contributions of tissue-level cellular composition and cell-autonomous transcriptomic maturation in predicting developmental age across diverse contexts.

**Figure 1.**
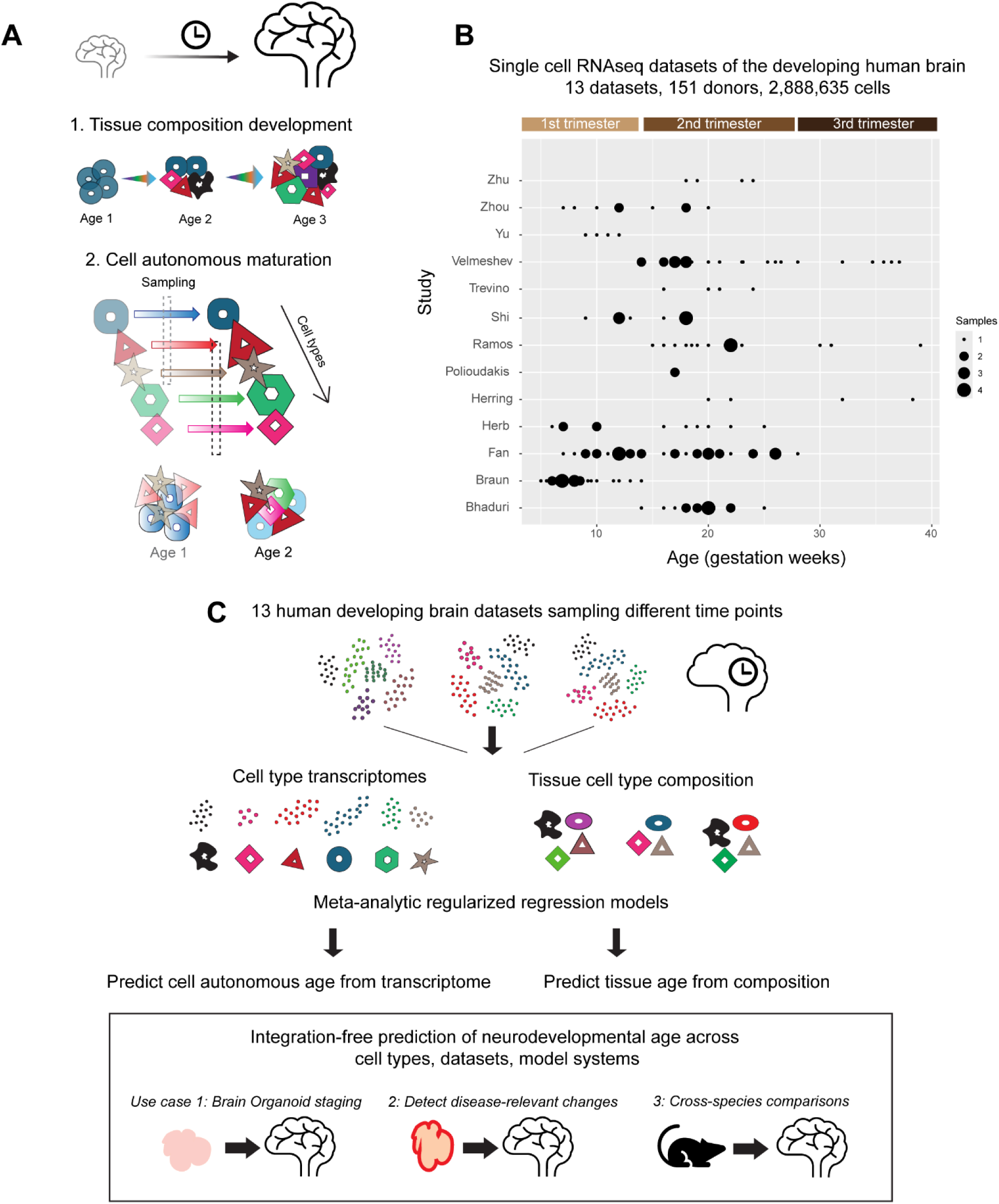
Meta-analysis of human brain single-cell data to identify tissue-level and cell-autonomous predictors of neurodevelopmental age. **A,** Neurodevelopment involves tissue-level changes in brain cell type composition and cell-autonomous transcriptomic maturation within individual cell types. Using a meta-analysis of recent single-cell datasets from the fetal human brain, we independently evaluate these features to identify robust patterns that consistently predict neurodevelopmental age across datasets. **B,** Distribution of sample ages in 13 fetal brain datasets included in meta-analysis. **C,** Summary of workflow: Meta-analytic regularized regression models are trained to identify cell autonomous or tissue-level predictors of neurodevelopmental age. We show their application for i) Assessing developmental progression in human neural organoids; ii) Identifying disease-associated shifts in mutant organoids, and iii) Aligning neurodevelopmental trajectories across human and mouse brain development.

### Tissue-level composition predicts neurodevelopmental age

#### Study-specific developmental changes in cell type composition do not generalize

Neurodevelopment is accompanied by large-scale shifts in cell type composition, following the well-established sequence of progenitor proliferation, neurogenesis, and subsequent gliogenesis ^41–43^. We first evaluated whether tissue-level cell type composition in the human brain reliably predicts developmental age across multiple datasets. To standardize comparisons, we grouped author-provided cell type annotations into seven broad cell classes based on consensus marker genes (**Fig 2A, Supplementary Fig 1**, **Supplementary table S3**) and trained regularized regression models to predict sample age from cell class proportions. Study-specific models performed well within individual datasets, accurately predicting gestational age in 7 of 11 studies, with strong correlations between predicted and actual ages (best performance: Braun, R = 0.94, *P* =2.6e-12; **Supplementary** Fig 2). Despite high within study-accuracy (median error = 1.19 weeks), these models failed to generalize across datasets, with prediction error increasing nearly 8-fold when applied to unseen studies (median error = 9.13 weeks, Wilcoxon test, *P* = 2.8*10^-6^; **Fig 2B**). Thus, while developmental changes in cell type composition are robust within individual studies, they do not generalize well across datasets.

**Figure 2.**
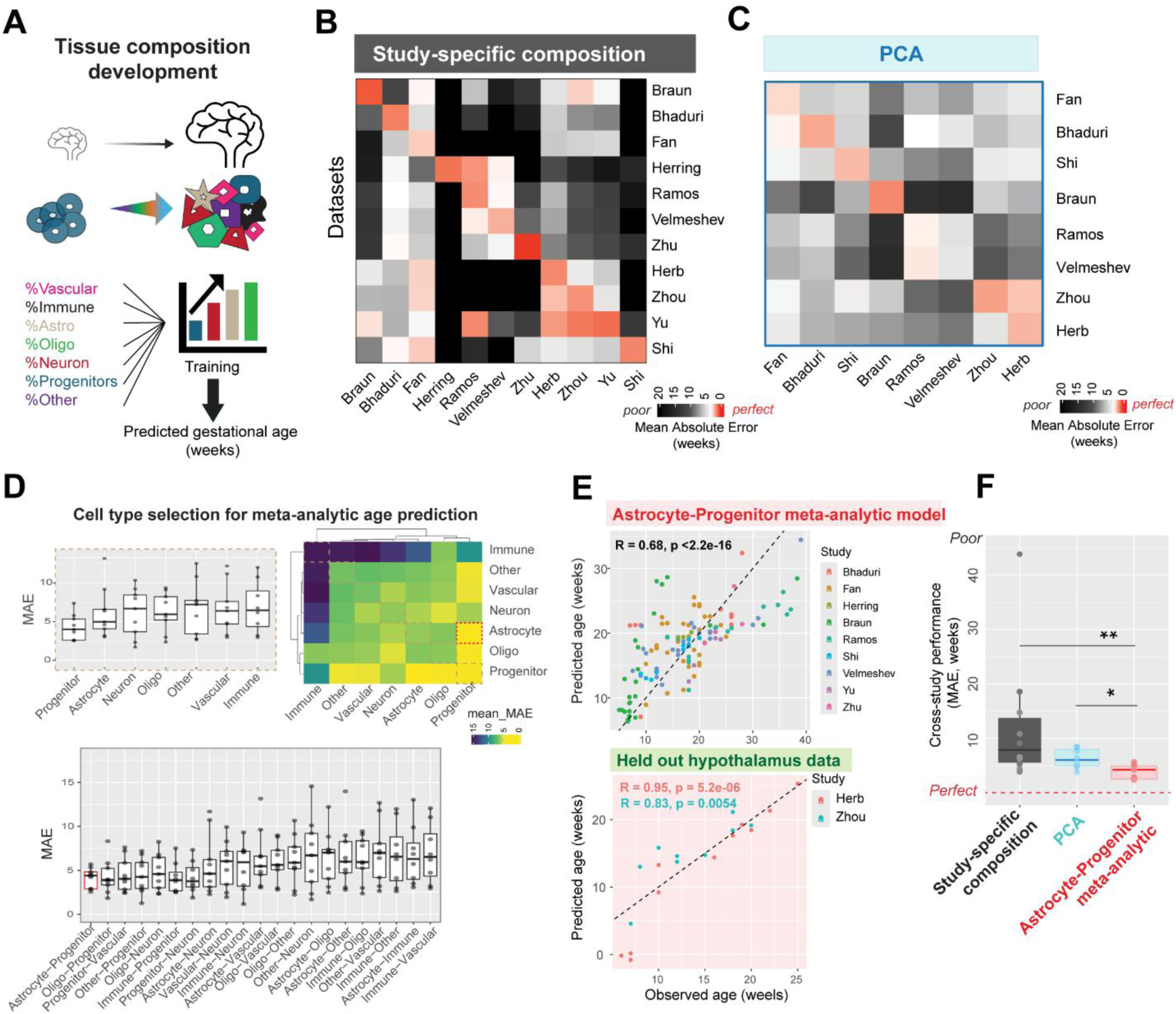
Tissue-level changes in cell type composition predict neurodevelopmental age, with study-specific differences impacting generalizability. **A,** Proportions of 7 broad cell classes-Vascular, Immune, Astrocyte, Oligodendrocyte, Neuron, Progenitor, and Others were used to train regularized regression models to predict sample age from tissue composition within individual studies. **B,** Within-study and cross-study performance (Mean Absolute Error in weeks) of study-specific compositional models (leave-sample-out cross validation, see Supplementary Fig 2). **C,** Performance of models trained on principal components (PCs): Linear models were trained on age-correlated PCs within each study derived from pseudo-bulked transcriptomes (see Supplementary Fig 3). The heatmap shows within-study and cross-study performance of PC-based age prediction. **D,** Linear models trained on proportions of specific cell types were evaluated for their ability to predict tissue age meta-analytically across datasets. Progenitors alone (top left) or the combination of progenitors and astrocytes (bottom) show least mean absolute error (MAE). The heatmap (top right) displays individual cell type performance along the diagonal and cell-type combination performance off the diagonal. **E,** Performance of meta-analytic model trained on astrocyte and progenitor proportions in leave-study-out cross validation (top). This model shows good performance in two held-out datasets from the human hypothalamus (bottom). **F,** Comparison of age-prediction performance (MAE) across held-out studies for three model types: study-specific composition, PCA, and astrocyte-progenitor proportions. The axis is restricted, with one value (792.63) from study-specific composition performance omitted for clarity. (\**P* < 0.05, \*\**P* < 0.01, Wilcoxon test).

### Principal component age predictions are primarily driven by cell type composition

Principal components (PCs) derived from gene-expression are commonly used to compare developmental age within a study, with PC1 typically showing strong correlation to age ^10–12,17^. However, because cell type composition is the dominant source of transcriptomic variation during development, it may underlie PC1-based age predictions. Given the poor generalizability of study-specific compositional changes, we assessed whether PC-based age predictions remain accurate across datasets using principal components obtained from pseudo-bulked transcriptomes.

In all eight studies tested, either PC1 (50%) or PC2 (50%) correlated significantly with age, enabling reliable within-study age prediction (median error = 1.95 weeks, **Supplementary** Fig 3). However, projecting other studies onto the age-correlated PC identified within a single study resulted in significantly worse performance, with a 3-fold increase in error (median error = 6.04 weeks, Wilcoxon test, *P* = 0.0003; **Fig 2C**). Since cell type composition varies across datasets, we examined the relationship between age-correlated PCs and cell type proportions. Strikingly, the correlation between PCs and age was nearly identical to their correlation with various cell type proportions (e.g., Astrocytes: R = 0.88, *P* = 2 * 10^-12^; Progenitor: R= −0.69, *P* = 1.1 * 10^-6^; Oligo: R = 0.81, *P* = 1.7 * 10^-10^; Immune: R = 0.78, *P* = 2.9 * 10^-9^; **Supplementary** Fig 3). Thus, principal components predominantly capture shifts in cell type composition to predict age within individual studies, but study-to-study variability in composition prevents PCs from reliably generalizing age predictions across datasets.

### Astrocyte and progenitor proportions consistently predict neurodevelopmental age across datasets

Rather than relying on global tissue composition, which varies across studies, we focused on well-defined cell types whose proportions are more consistent and reliably predict developmental age. Using feature selection, we identified the best-performing cell classes for age prediction across datasets. Linear models using only progenitors, or a combination of progenitors and astrocytes, achieved the best performance across studies in a leave-study-out cross validation (Mean error across cross validation folds, using progenitors: 4.26 weeks, using astrocytes and progenitors: 4.14 weeks, **Fig 2D**). The meta-analytic model trained using astrocyte and progenitor cell class proportions reliably predicts age across 9 studies (correlation between predicted and actual age of samples: R = 0.68, *P* < 2.2 * 10^-16^, **Fig 2E**). We further tested this pre-trained model on two held-out datasets from the developing human hypothalamus ^15,33^, achieving accurate predictions of age in both studies (correlation between predicted and actual ages: Herb: R = 0.95, *P* = 5.2 * 10^-6^; Zhou: R= 0.83, *P* = 0.005; **Fig 2E**).

A simple compositional model using astrocyte and progenitor cell class proportions effectively predicts age in held out datasets with least error (median error = 4.52 weeks), compared to study-specific composition (median error = 9.13 weeks; Wilcoxon test: *P* = 0.001), or PCA (median error = 6.03 weeks; Wilcoxon test: *P* = 0.007; **Fig 2F**). This model relies on the increase in astrocyte proportions and the decrease in progenitor proportions during development, depicted with a positive coefficient for astrocytes (35.32) and negative coefficient for progenitors (−22.46).

In sum, study-specific tissue composition predicts neurodevelopmental age within individual studies, but cross-study variability prevents this from generalizing across datasets. In contrast, astrocyte and progenitor proportions exhibit consistent developmental trajectories across datasets, enabling robust and generalizable age predictions. These relative shifts in astrocyte and progenitor abundance are sufficient to predict tissue age across studies.

We next turned our attention to cell autonomous predictors of developmental age trained on single-cell transcriptomes that are independent of tissue composition.

### Cell type-agnostic transcriptomic signatures enable reliable predictions of cell autonomous neurodevelopmental stage

Single cell transcriptomic age predictors have enabled reliable predictions of chronological age in aging tissues from mice ^44^ and humans^45^. We aimed to develop a transcriptomic age predictor for cell types in early human brain development to uniformly compare cell autonomous maturation across datasets. To mitigate the impact of single-cell data sparsity, we used log normalized expression from cell type aggregated transcriptomes to train meta-analytic regularized regression models. Models were first trained on each specific cell type to predict age in held-out datasets (leave-study-out cross validation, **Fig 3A**). Cell type-specific models reliably predicted the age of the cell class they were trained on across datasets (Median error across cross validation folds in weeks: Astrocyte: 4.10, Progenitor: 2.924, Excitatory Neuron: 2.27, Inhibitory neuron: 2.31, Oligodendrocytes: 3.82, Vascular: 5.86, Immune: 4.05, Other: 4.58; **Fig 3B**).

**Figure 3.**
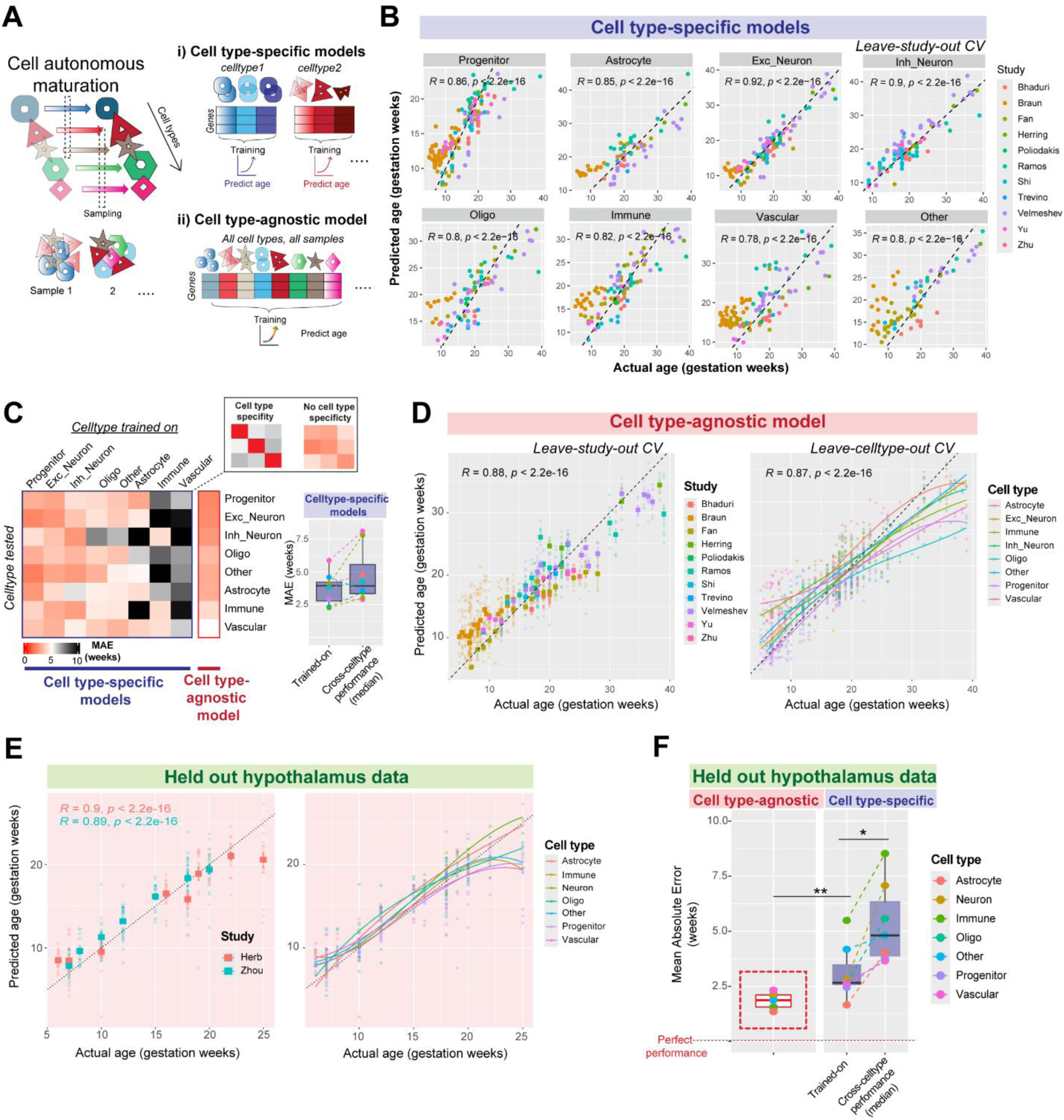
A cell type-agnostic model robustly predicts neurodevelopmental age across cell types and datasets. **A,** Cell-autonomous transcriptomic maturation occurs within individual cell types during neurodevelopment and distinct samples across datasets may capture cells in different states of maturation. We trained meta-analytic cell type-specific models on individual cell types, or a cell type-agnostic model jointly on all cell types to predict age from log-normalized gene expression. **B,** Performance of cell type-specific models in predicting neurodevelopmental age of cell types from held-out studies (leave study out cross validation). **C**, Heatmap shows cross-cell type performance of cell type specific models (left), revealing poor cell type specificity, akin to the performance of a cell type-agnostic model. The boxplot (right) compares the performance of each cell type-specific model on its trained cell type with median performance across all other cell types. **D,** Performance of cell type-agnostic model on predicting age across held out studies (leave-study out cross validation, left) and across held out-cell types (leave-cell type out cross validation, right). **E,** Cell type agnostic model shows robust performance in two held out hypothalamus datasets not used in training. **F,** Comparison of mean absolute error (MAE) in age prediction within held out hypothalamus datasets by the cell type-agnostic model (left) and cell type-specific models (right, within and across cell types; \**P* < 0.05, \*\**P* < 0.01, Wilcoxon test).

However, cell type-specific models also showed equivalent age prediction accuracy in cell types that they were not trained on, revealing a surprising lack of cell type specificity (median error on trained-on cell types: 3.94 weeks, median error on other cell types: 3.92 weeks; Wilcoxon test: *P* = 0.25, **Fig 3C**).

Given the lack of cell type specificity observed above, we deliberately trained a cell type-agnostic model on transcriptomes from all cell classes across datasets to predict age. By equally weighing every cell type within each study, this cell type-agnostic model learned a general signature of neurodevelopment, showing robust performance in predicting age across held out studies (median error: 2.65 weeks), and held out cell types (median error: 2.86 weeks; **Fig 3D**). To further test the generalizability of this model, we applied it to two held-out hypothalamus datasets ^15,33^, obtaining accurate predictions in both (Correlation between actual and predicted ages, Herb: R = 0.9; Zhou: R = 0.89; **Fig 3E**). The cell type agnostic model achieved the best performance across all cell types in held out hypothalamus data (median error: 1.87 weeks), outperforming cell type-specific models (median error: 2.65 weeks, Wilcoxon test: *P* = 0.0069, **Fig 3F**).

The robustness of this model comes from a diverse training dataset that broadly samples cross-cell type and cross-study variability. In contrast, models trained on data from only one study show good performance within that study (median error: 1.52 weeks) but fail to generalize to other studies (median error: 8.03 weeks, Wilcoxon test: *P* = 0.0078). In sum, a meta-analytic cell type agnostic model robustly predicts age across cell types and datasets of the developing human brain, eliminating the need for rigorous cell type alignment across datasets. Next, we carefully evaluated features used by the model, identifying gene sets that contribute to its robust performance.

### Sampling diverse neurodevelopmental programs drives robust predictions of cell type agnostic model

The regularized cell type agnostic model weighted only 462 genes, a small fraction of the transcriptome, to robustly predict developmental age across cell types and datasets. Among these, 191 genes have negative coefficients, and 271 have positive coefficients (top 20 shown in **Fig 4A; Supplementary table S4**). The average expression of genes with positive coefficients correlated positively with gestational age (R = 0.83), while genes with negative coefficients was negatively correlated to age (R = −0.44, **Fig 4B**). We identified age-correlated genes within each study (**Supplementary table S5**) and compared them to the coefficients assigned by the meta-analytic model. Genes with a positive coefficient generally show positive age correlation in all studies, while genes with negative coefficients tend to be negatively correlated to age. In the classification task of distinguishing positive and negative model genes based on study-specific age correlations, the average AUROC performance across studies is 0.74 (**Fig 4C**). Notably, the model-selected genes are **not** the most strongly age-correlated genes individually: A classifier trained on absolute correlation coefficients fails to distinguish them from the rest of the genome (AUROC = 0.54).

**Figure 4.**
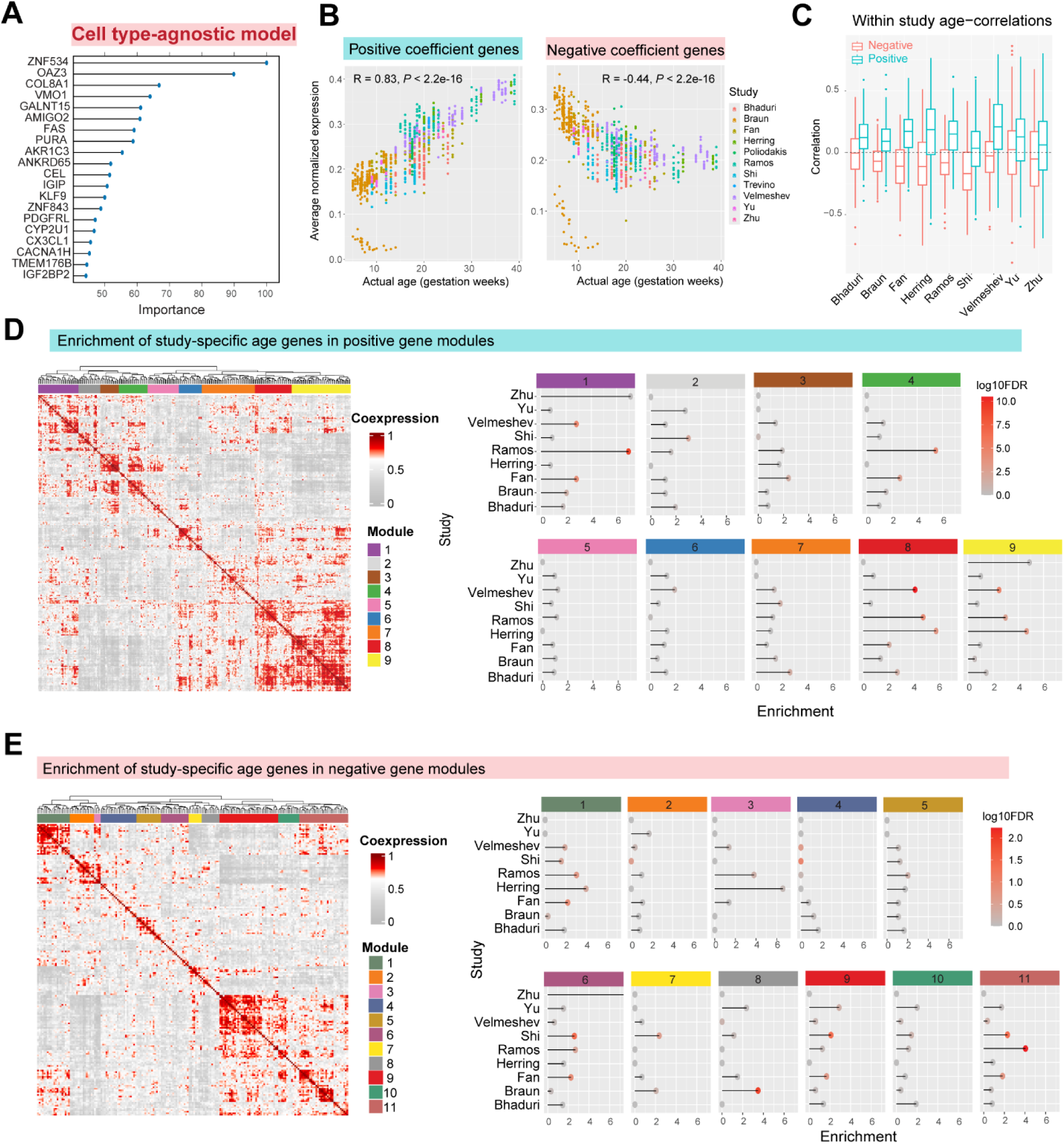
Co-expression analysis of cell type-agnostic model genes reveals distinct contributions of study-specific developmental programs. **A,** Top 20 important genes by absolute coefficient in the cell type-agnostic model. **B,** Average expression of positive coefficient genes (left) and negative coefficient genes (right) across all datasets. **C,** Boxplot shows correlation coefficients of model genes with age within individual studies. Positive genes generally show higher correlation coefficients within every study compared to negative genes (*P* < 10^-3^ for all studies, Wilcoxon test with FDR correction). **D-E,** Aggregate co-expression network of positive (D) and negative model genes (E) hierarchically clustered to identify distinct co-expression modules (9 positive and 11 negative modules). Panels on right show enrichment of study-specific age correlated genes within each module. Each module represents age-genes from distinct study combinations.

Overall, the cell type-agnostic model captures the directionality of age-related expression changes within individual studies, even though the selected genes are not the strongest individual markers of developmental age. Instead, their aggregate expression encodes a robust developmental signal preserved across cell types and datasets. To understand the biological basis of this predictive power, we examined the representation of distinct biological processes and study-specific age genes in the model.

First, we clustered positive and negative coefficient model genes into co-expression modules and then evaluated the enrichment of study-specific age genes in each module (**Fig 4D-E, Supplementary table S6**). Our analysis revealed that each module consists of age-correlated genes from distinct studies, that represent diverse biological processes (**Fig 5, Supplementary table S7**). For example, among positive gene modules (**Fig 4D**), Module 4 is highly enriched for age genes from Ramos and Fan datasets and represents amino acid transporter activity. Modules 8 and 9 are both highly enriched for genes from Velmeshev, Ramos, and Herring datasets, and represent glial cell proliferation and synaptic transmission respectively. Modules 5 and 6 are not enriched for any one dataset but represent lipid catabolism and action potential-related genes respectively (**Fig 5A**). Similarly, among negative gene modules (**Fig 4E**), Module 9 is enriched for Shi, Yu, and Fan datasets and represents neuropeptide signaling and GPCR activity, while Module 11 is enriched for Shi, Ramos, and Fan datasets and represents cholesterol metabolism. Module 8 is specifically enriched for the Braun dataset and represents postsynaptic cytoskeletal organization (**Fig 5B**).

**Figure 5.**
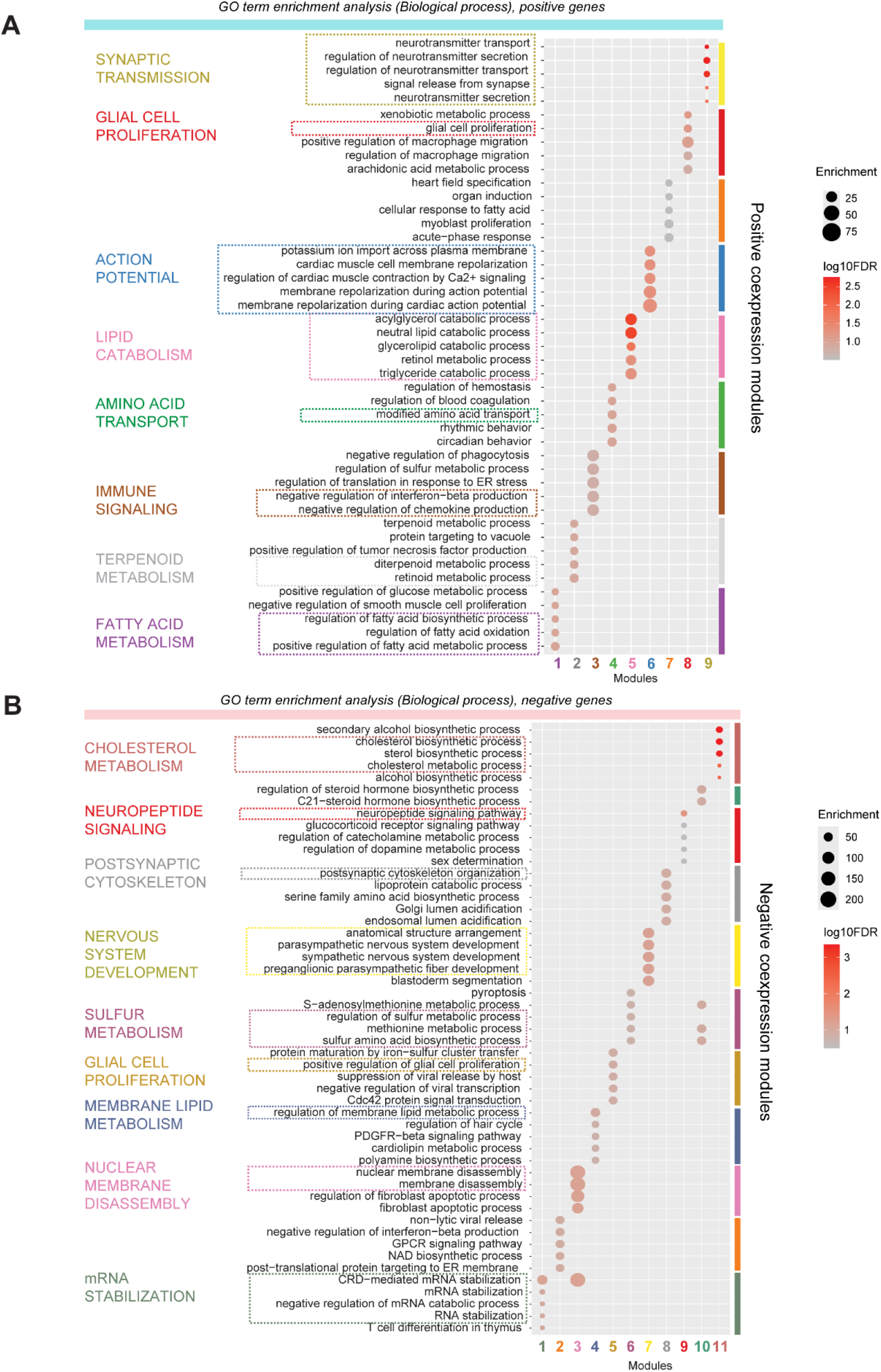
GO enrichment of cell type-agnostic model genes highlights diverse biological processes contributing to developmental age prediction. **A,** Positive gene modules are significantly enriched for distinct GO biological process terms including synaptic transmission, action potential, and lipid catabolism. **B,** Negative gene modules are significantly enriched for cholesterol metabolism, postsynaptic cytoskeleton, nervous system development etc.

In sum, genes used by the cell type-agnostic model form distinct co-expression modules enriched for age-associated genes from different studies. Regularization selects genes that may not be individually predictive but collectively ensure robust performance, capturing diverse biological processes essential for neurodevelopment.

As an alternate approach to characterize gene sets driving model performance, we individually screened thousands of GO terms by training models exclusively on genes from a single GO term. We expected GO term models to generally perform worse than models using broader gene sets but anticipated few GO terms that would excel in a cell type-specific manner. Totally, 2213 GO term-based models, each with 50 to 500 genes were trained (**Fig 6A**). Models trained on synGO ^46^, a set of 1112 synapse-related genes were also used for comparison. As expected, only a minority of GO term models showed better performance than models using all genes: in astrocytes: 30.6 %, vascular cells: 19.1 %, progenitors: 17.1 %, oligodendrocytes: 13.1%, other: 8.1 %, inhibitory neurons: 0.99 %, excitatory neurons: 0.54 %, and immune cells: 0.49 %. We identified top-performing GO terms per cell type (**Fig 6B, Supplementary table S8**), which differ across cell types resulting in poor cross-cell type correlations (Spearman correlation coefficient < 0.5, **Fig 6 C-D**). Ultimately, the highest accuracy in age-prediction was achieved by the cell type-agnostic model trained on all genes rather than any individual GO term (empirical p-value = 0.00045; **Fig 6A**).

**Figure 6.**
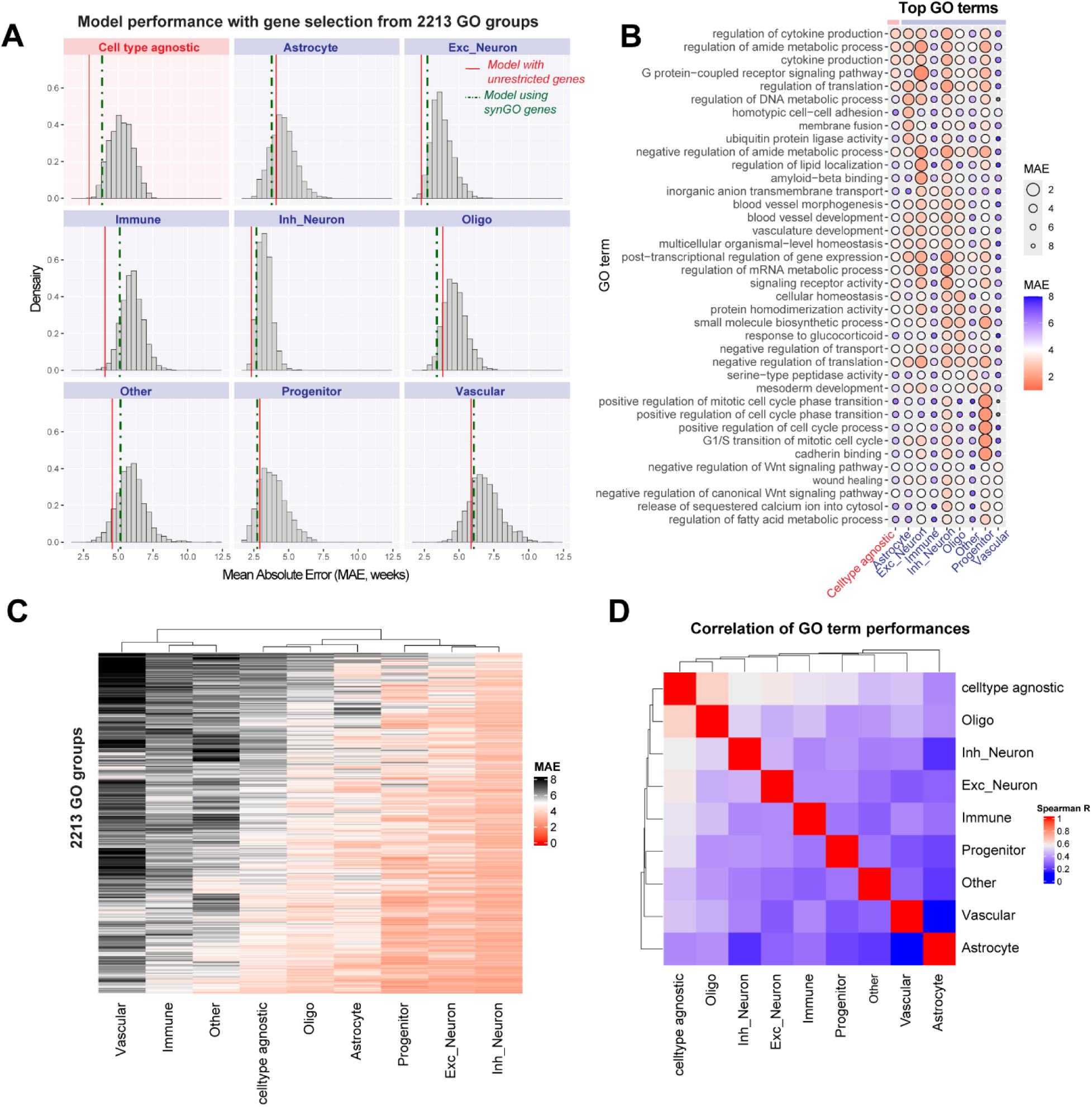
Characterization of GO terms driving age prediction performance reveals cell type-specific signatures. **A,** Age prediction models were trained using restricted gene sets derived from GO groups with 50-500 genes. Distribution of performances is shown for each of the cell type-specific and cell type agnostic models. For comparison, red vertical lines indicate the performance of models trained using all genes (from Fig 3), and dashed vertical lines indicate performance of models using the synGO gene set. **B,** Dotplot shows the performance of top 5 best-performing GO terms in each cell type. In astrocytes: regulation of DNA metabolic process; progenitors: positive regulation of mitotic cell cycle phase transition; excitatory neurons: GPCR signaling pathway; inhibitory neurons: regulation of translation; oligodendrocytes: cellular homeostasis; immune cells: inorganic anion transmembrane transport; vascular cells: negative regulation of Wnt signaling pathway. **C,** Heatmap shows performance of all GO terms in each cell type. **D,** Spearman correlation of performance across all GO terms between cell types.

These findings demonstrate that multiple distinct biological processes drive developmental age prediction, highlighting the robustness and degeneracy in molecular clocks of human neurodevelopment. Various strategies can be used to prioritize gene sets for either cell type-specific or cell type-agnostic age predictions. Cell type-specific accuracy is improved by focusing on biological processes relevant to that cell type, while robust cross-cell type predictions are achieved by sampling diverse biological processes.

Using our suite of developmental age predictors, we next probed whether these signals are preserved in other model systems of neurodevelopment. Specifically, we asked whether cell types in human neural organoids cultured *in vitro* recapitulate the tempo of cell-autonomous maturation seen *in vivo*.

### Cell type-agnostic model trained on primary tissue predicts developmental progression in human neural organoids

Human neural organoids are self-organizing 3D cellular aggregates derived from pluripotent stem cells, with the potential to recapitulate the cellular diversity of the developing human brain ^38,47–49^. Organoids enable the characterization and manipulation of cell types that are otherwise inaccessible. While individual cell types in organoids are thought to align closely to the human brain ^20,50,51^, the extent to which lab-grown organoids reliably recapitulate the tempo of *in vivo* neurodevelopment is a topic of ongoing research ^11,29,52^. High variability across batches, cell lines, and protocols has been reported previously ^53^. We aimed to determine the developmental age of cell types in organoids with respect to the human brain to enable uniform comparisons.

Using our cell type-agnostic model (Fig 3), we assess developmental age in organoid cells across diverse protocols (**Fig 7**) without needing to integrate or align cell type annotations across datasets. The cell type-agnostic model trained on human fetal cell types accurately predicts normal developmental progression in organoid cell types (**Fig 7A**), supporting the preservation of *in vivo* molecular programs in cultured organoids. Predicted age across all cell types is highly correlated to *in vitro* culture age in both cortical (R = 0.91, **Fig 7A**) and midbrain organoids (R = 0.89) from Uzquiano et al. 2022, and Fiorenzano et al. 2022, respectively ^20,39^. Furthermore, in a dataset where cultured organoids were transplanted into the rat brain ^40^, our model accurately predicts accelerated maturation in transplanted organoids (mean age = 32 weeks) compared to non-transplanted organoids (mean age = 24 weeks, *P* < 2.2 * 10^-16^).

**Figure 7.**
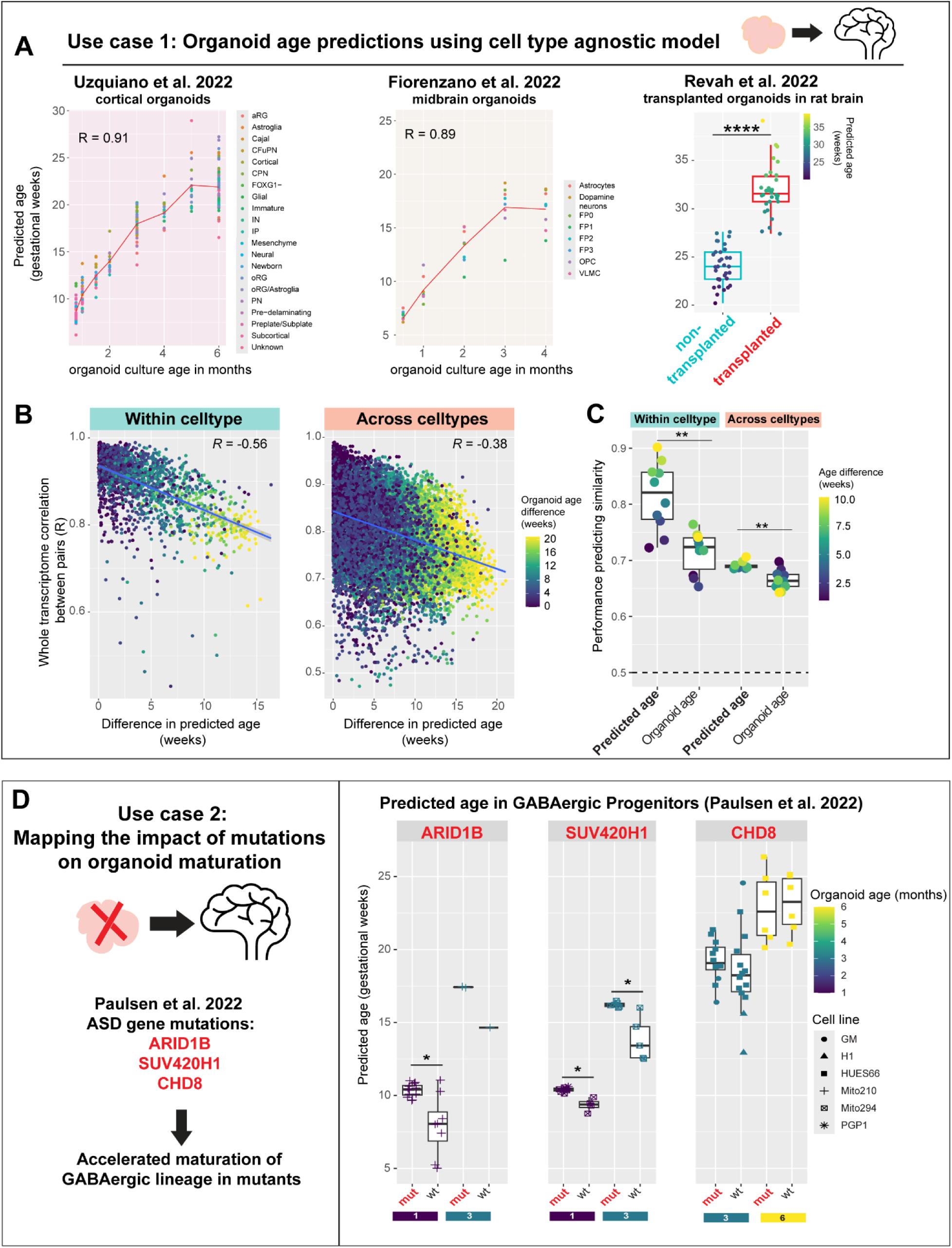
Cell type-agnostic model predicts normal developmental trajectories and disease-associated shifts in human neural organoids. **A,** Performance of the cell type agnostic model (from Fig 3) in predicting developmental age in human neural organoid datasets from three different protocols. Predicted ages are highly correlated to the in vitro culture age of organoids in cortical (left) and midbrain organoids (middle). Cortical organoids transplanted into the rat brain show higher predicted ages compared to non-transplanted organoids (right, \*\*\*\**P* < 10^-4^, Wilcoxon test). **B,** Whole-transcriptome similarity of cell pairs (Pearson’s R) is negatively correlated to their difference in predicted age both within and across cell types. **C,** Performance in predicting cell-cell similarity based on predicted age vs observed organoid age (days in vitro). \*\**P* < 0.01, Wilcoxon test. **D,** Developmental age prediction in cortical organoids with mutations in ASD-linked genes: ARID1B, SUV420H1, and CHD8. Right: Predicted age of GABAergic progenitor cell types is significantly elevated in ARID1B and SUV420H1 organoids at both 1 and 3 months of culture (\**P* < 0.05, Wilcoxon test), matching predictions from the original study. CHD8 mutants show a minor increase in 3-month-old organoids.

Independently, we assessed the relevance of predicted age in organoid cells by examining its ability to predict whole transcriptome similarity. We expected that cells closer in predicted age would be more correlated than cells with different predicted ages. Correlation between cells was indeed negatively correlated to the difference in their predicted ages, both within (R = −0.56) and across cell types (R = −0.38; **Fig 7B**). We further used an AUROC-style metric (**Fig 7C**) to assess the correlation between cell pairs of similar ages (e.g. age difference < 3 weeks) versus different ages (e.g. age difference > 3 weeks). This metric consistently shows higher performance when computed using predicted age rather than the actual organoid culture age. For instance, comparing cell pairs with age difference greater or lesser than 3 weeks, AUROC = 0.77 with predicted age versus 0.65 with organoid culture age. Overall, predicted ages from the cell type-agnostic model show significantly higher performance in predicting cell-cell similarity compared to actual organoid culture age both within (*P* = 0.0022) and across cell types (*P* = 0.0017, Wilcoxon test).

We assessed whether our model detects cell type-specific temporal shifts induced by disease-causing mutations, using cortical organoids with autism spectrum disorder (ASD)-linked mutations ^54^ (**Fig 7C**). Strikingly, the model predicts significantly older ages specifically in GABAergic progenitors of ARID1B mutants relative to wild type organoids at both 1 and 3 months of culture (mutant - wildtype age difference at 1 month: 2.40 weeks, *P* = 0.016; at 3 months: 2.76 weeks, n=1). A similar pattern is also detected in SUV420H1 mutants (mutant - wildtype age difference at 1 month: 1.05 weeks, *P* = 0.0040; at 3 months: 2.37 weeks, *P* = 0.0086). In CHD8 mutants, we observe slightly older GABAergic progenitors in 3-month-old organoids from the HUES66 cell line (mutant - wildtype age difference: 0.88 weeks). These results align with the accelerated maturation of the GABAergic lineage in mutant organoids reported in the original study, which found a shift in the pseudotime trajectories of mutant GABAergic cells. Our model independently provides quantitative estimates for this age-shift in each cell type. Age predictions per cell line, culture age, and lineage for all organoids are shown in **Supplementary figures 4-6**.

We further evaluated the robustness of our model in the Human Neural Organoid Cell Atlas (HNOCA), an integrated meta-atlas of 36 organoid single-cell datasets with 1.77 million cells from multiple protocols ^27^. A challenge with large integrated datasets is the use of different technologies across studies, resulting in varying read depths that could affect age-prediction from log-normalized expression. Therefore, we used a normalization-robust model variant that uses rank normalized expression (see methods). This model successfully predicts developmental progression of cell types across all studies in the HNOCA atlas (**Supplementary** Fig 7), with overall correlation coefficient of 0.52 between the predicted age and organoid culture age.

Consistent with known metabolic limitations of organoids grown in culture long-term ^11^, predicted ages decline after about 6 months in culture (**Supplementary** Fig 7B). Predicted age is most advanced by transplantation *in vivo*, as the oldest cell types in our organoid analysis are present in the rat-transplanted organoids from Revah et al. 2022.

These results establish the generalizability of cell type-agnostic age predictors across diverse human neural organoid datasets. Our method enables uniform comparisons across organoid protocols, reveals lineage-specific temporal shifts caused by mutations, and extends to large integrated atlases. Next, we investigated whether models trained on human data could generalize to the embryonic mouse brain, aiming to assess evolutionary conservation.

### Models trained on human brain tissue accurately predict accelerated pace of embryonic mouse brain development

The human brain develops at a slower pace compared to other species, a phenomenon called neoteny or bradychrony ^55,56^, whereas mice are thought to exhibit accelerated neurodevelopment. The mechanisms underlying differences in developmental tempo across species remain largely unknown. We aimed to compare cell-autonomous maturation across human and mouse neurodevelopment by using age predictors trained on human data, restricting to genes with mouse orthologs.

First, we assessed whether cell-autonomous programs of human neurodevelopment were conserved in mice by evaluating the performance of a model trained purely on human data in an embryonic mouse brain dataset ^57^. Strikingly, the cell type-agnostic model trained on human tissue accurately predicts developmental progression of cell types in the mouse brain (**Fig 8A**). Predicted human age is strongly correlated to the actual embryonic mouse age in neurons (R = 0.94), neuroblasts (R = 0.94), glioblasts (R = 0.9), radial glia (R = 0.9), Vascular cells (R =0.94), oligodendrocytes (R = 0.76), and immune cells (R = 0.47). However, very early non-neural cell types absent in the human training data show poor performance, including endoderm (R = −0.5) and mesoderm cells (R = 0.0084). Intriguingly, we observe a transient decrease in predicted age around E8.5 in mouse neural progenitor cells, which might correspond to a similar “rejuvenation event” observed in the epigenetic DNA methylation age of mouse embryonic stem cells ^58^.

**Figure 8.**
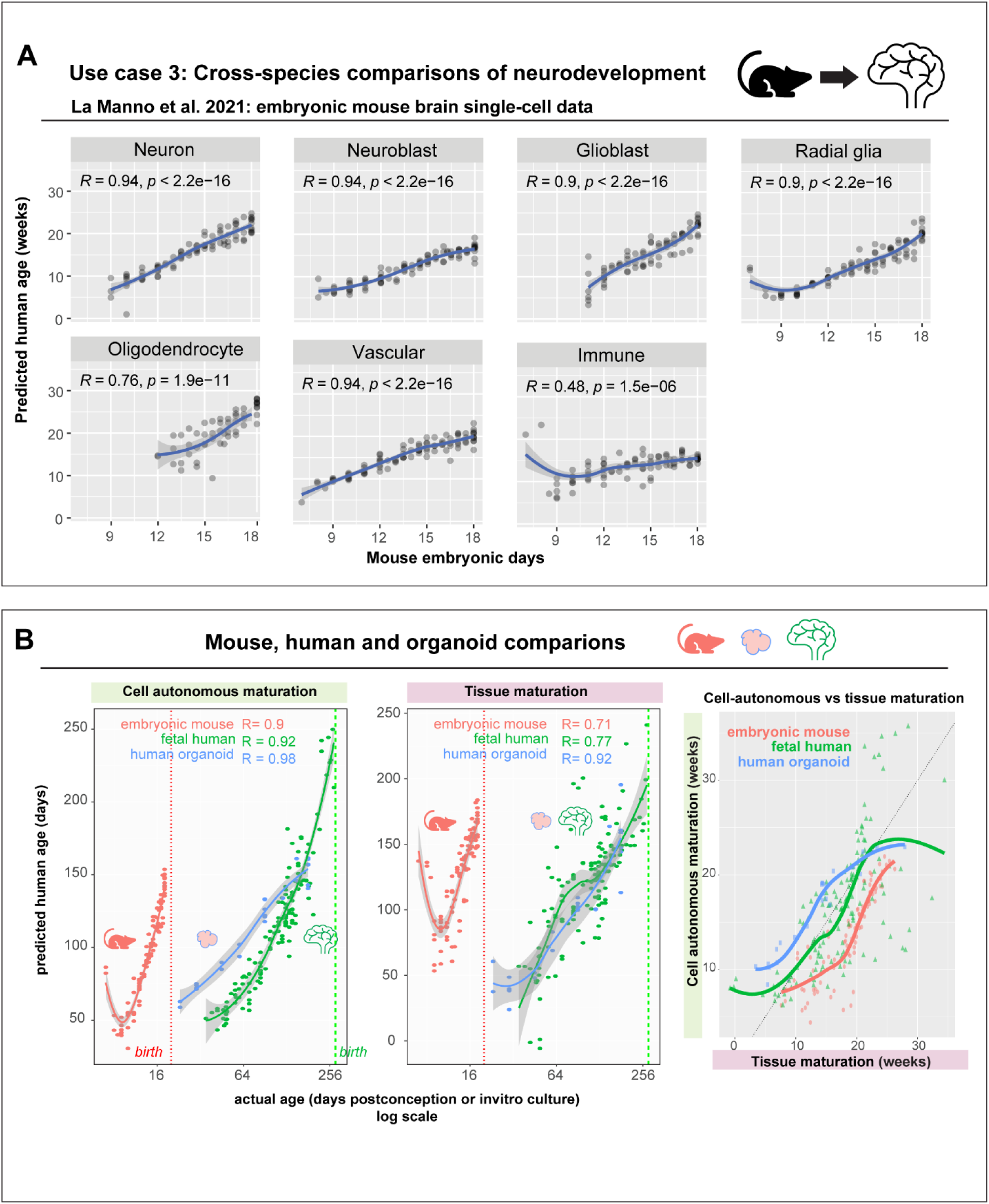
Cross-species generalization of model trained on human brains enables uniform comparisons of neurodevelopment across mice, human brains and organoids. **A,** Performance in predicting developmental age of cell types (in human weeks) in embryonic mouse brain development using a cell type-agnostic model trained on human data. Genes were restricted to mouse orthologs**. B,** Comparing predicted cell-autonomous ages across mouse, human brain development, and human neural organoids (left). Middle panel shows tissue compositional age predicted from proportions of astrocytes and progenitors per sample in mouse, human brains, and human neural organoids. Average predicted cell-autonomous age of neurons, glia, and progenitors per sample are plotted against the compositional age, revealing a synchrony between cell-autonomous maturation and global tissue maturation across human, mouse brains and organoids.

Next, we uniformly compared cell autonomous maturation across the developmental span of human, mouse brains, and human neural organoids. By comparing the average predicted age of neurons, astrocytes and progenitors per sample against actual age (**Fig 8B**), we observe accelerated cell autonomous maturation in mice compared to human brains and neural organoids (slope: mice = 9.762, humans = 0.811, human neural organoids = 0.550), with mice reaching maturation equivalent to 21 human weeks in just 18 days (ANOVA, effect of tissue: F _(2,249)_ = 260.59, *P* < 2e-16). We further compared the tempo of global tissue-level development using meta-analytic models based on astrocyte and progenitor proportions from Fig 2. This compositional model accurately predicts developmental progression in organoids and mice brains, recapitulating patterns seen with cell autonomous maturation (slope: mice = 8.04, humans = 0.74, organoids = 0.58). The tempo of cell-autonomous maturation and global tissue-level development appear synchronized across human brains, mouse brains and organoids with no tissue-specific differences (ANOVA, interaction between compositional age and tissue, F_(2,249)_ = 0.765, *P* = 0.466). While these results require validation in additional datasets and species, they provide a framework to evaluate conserved, coordinated neurodevelopmental changes across model systems.

Our work establishes a foundation for uniformly comparing neurodevelopmental age across diverse contexts, model systems, and species. We provide robust integration-free, cell type-agnostic age prediction models that will allow users to assess neurodevelopmental progression in their own single-cell datasets.

## Discussion

We analyzed 13 human fetal brain datasets with over 2.8 million cells to identify robust tissue-level and cell-autonomous predictors of neurodevelopmental age. We show that specific cell type proportions reliably predict tissue age across datasets and further train a cell type-agnostic model to track cell autonomous maturation in diverse cell types. This model generalizes across multiple human brain, human neural organoid and mouse brain datasets, revealing disease-specific and evolutionary shifts in neurodevelopmental tempo.

While individual studies have successfully measured developmental progression within their single-cell datasets ^11,19,28^, identifying robust patterns that generalize across multiple studies is more challenging. Cross-study variability in cell type composition obscures tissue-level developmental changes, and known batch effects in single-cell data ^59^ further challenge the identification of true cell-autonomous features of neurodevelopment. Encouragingly, meta-analytic models trained on diverse datasets can extract highly reproducible tissue-level and cell autonomous predictors of developmental age. Accounting for substantial cross-study variability enables robust developmental age prediction across human brain and neural organoid datasets, and most strikingly, demonstrates cross-species robustness.

A major advantage of our work is the prediction of age across diverse cell types using a cell type-agnostic model. This is particularly relevant for stem cell-derived human organoid models where recent efforts have focused extensively on annotating *in vitro* cell types ^27,60,61^. Comparing developmental stage of cells across diverse organoid protocols has remained challenging due to the difficulty in matching cell types across protocols. By using a pre-trained cell type-agnostic model, we directly predict age in cell types across diverse neural organoid protocols, including the integrated human neural organoid cell atlas (HNOCA). This has the potential to identify interventions that alter maturation rates *in vitro*, as demonstrated by our prediction of lineage-specific shifts in organoids with ASD mutations ^54^

A striking feature of the cell type-agnostic model is its ability to predict developmental progression in the embryonic mouse brain, despite being trained only on human brain tissue. Comparing predicted ages across human and mouse gestational span reveals greatly accelerated maturation during mouse brain development, in line with the known neoteny of human neurodevelopment ^62,63^. In predicting developmental age from the cell’s transcriptome, our models are also sensitive to the influence of the extracellular environment on developmental tempo. Notably, we accurately detect accelerated maturation in human neural organoids transplanted into the rat brain. *In vitro* models are a major avenue for studying cross-species differences in developmental tempo ^64–67^, and the extent to which the pace of development is set by cell intrinsic or extrinsic factors is a topic of ongoing research. Our framework to compare predicted tissue compositional age and cell autonomous age across species and organoid models may prove useful for quantitative assessment of inter-species differences ^68^

We establish meta-analysis as an effective strategy for identifying conserved signatures in early neurodevelopment. This approach enables standardized comparisons of neurodevelopment from single-cell data, complementing previous studies on late-life aging ^44,45,69–71^. While incorporating additional datasets could further enhance robustness, our meta-analytic models provide a powerful framework for comparing developing brain cells across contexts, model systems, and species. These findings pave the way for optimizing *in vitro* models of neurodevelopment, uncovering disease mechanisms and advancing therapeutic interventions.

## Data and code availability

This study did not generate new unique reagents. This paper analyzes existing, publicly available data, accessible at links provided in Supplementary table S2. All original code used for analysis will be made available at the time of publication at https://github.com/sridevi96/NeuroDevTime. Any additional information required to reanalyze the data reported in this paper is available from the lead contact upon request.

## Acknowledgements

We would like to thank Dr. Leon French, Dr. Hamsini Suresh, and Dr. Julien Muffat for helpful discussions. This research was generously funded by NIH U24MH130968 (JG, JMW), NIH R01MH133181 (JG), NIH R01MH113005 (JG), CIHR PJT-180565 (YL), and Schmidt Science Fellows (SV).

## Author contributions

SV, JG and YL conceptualized the experiments, SV and JMW curated the datasets; SV and JG conducted the analysis; SV wrote the manuscript with revision and editing by JG and YL.

## Supplemental information

Document S1. Supplementary Figures 1–7

Table S2. Excel file with information on datasets used for meta-analysis.

Table S3. Excel file with cell type mapping.

Table S4. Excel file with coefficients of cell type agnostic model, related to Fig 3.

Table S5. Excel file with list of study-specific age correlated genes, related to Fig 4.

Table S6. Excel file with co-expression modules derived from model genes related to Fig 4.

Table S7. Excel file with GO enrichment results of co-expression modules related to Fig 5.

Table S8. Excel file with performances of GO term models related to Fig 6.

## STAR Methods

### Dataset download and pre-processing

Links for all downloaded datasets used in our meta-analysis are provided in **Supplementary table S2**. A total of 13 human fetal brain datasets were used. Analysis was conducted using Seurat (v 5.1). Processed seurat objects for 7 human fetal brain datasets ^30–33,35–37^ and 2 human neural organoid datasets ^20,40^ were sourced from our previous meta-analysis ^52^. For other datasets, processed data provided by authors were used directly without additional filtering. Raw expression counts were log-normalized using Seurat’s NormalizeData (normalization.method = “LogNormalize”, scale.factor = 10000) and used for all analyses unless otherwise specified. Genes were restricted to common genes found across all human fetal brain and neural organoid datasets resulting in a final set of 10,957 genes.

Author-provided metadata from the original publications were used to group cells by batch, age, and cell type. We mapped the author-assigned cell types to 7 broad cell types (Astrocyte, Progenitor, Neuron, Oligodendrocyte, Immune, Vascular, Others; **Supplementary table S3**). We validated our cell type groupings by estimating overlap of top 100 consensus marker genes per cell type across datasets (**Supplementary** Fig 1). Consensus markers for each author-assigned cell type were identified as recurrent marker genes across all samples in the study using Seurat’s FindAllMarkers function, with criteria of log2FC > 2 and pct.1 > 0.2.

### Compositional age prediction models

#### Study-specific compositional models

Generalized linear models were trained to predict age (gestation weeks) of each sample within a study from the total proportions of each cell type present in that sample. Models were trained using Elasticnet regression (method = “glmnet”) from the caret package (6.0.94) with leave one sample out cross validation (trainControl method = “loocv”). Only studies with at least 4 different time points were used for this, resulting in a total of 11 study-specific compositional models (**Supplementary** Fig 2). Mean Absolute Error (MAE) was used to compare performance of each model within and across studies. Correlation coefficients and p-values shown are from the stat_cor function in ggpubr.

#### Principal Component Analysis

Raw expression counts within each sample were pseudo-bulked using Seurat’s AggregateExpression (group.by = “orig.ident”). Principal components were computed from pseudo-bulked expression matrices using Seurat’s RunPCA (npcs = 5) within each dataset with more than 4 distinct time points. PC1 or PC2 were significantly correlated to sample age in each dataset (**Supplementary** Fig 3). A generalized linear model (GLM) was trained within each study to predict age using the principal component (PC) most strongly correlated with age, resulting in a total of 8 study-specific models. Each study-specific model was tested on other datasets by projecting them onto the age-correlated PC used by the model and predicting age with the GLM (Fig 2C). MAE was used to compare performance of each model within and across studies.

#### Meta-analytic compositional model

We performed feature selection to identify cell types whose proportions reliably predicted age across studies. Meta-analytic linear models (method = “lm”) were trained on 9 datasets to predict age in each held out dataset (leave study out cross validation) using the proportions of individual cell types or combinations of 2 cell types. The best performing cell type combination, astrocytes and progenitors, was identified to show least MAE across held out datasets. Robustness of this astrocyte-progenitor meta-analytic model was tested further in two datasets (Herb and Zhou) that were held out during training.

### Cell autonomous age prediction models

#### Cell type-agnostic model

Single cell transcriptomes were aggregated by author-assigned cell type within each sample in a dataset using Seurat’s AggregateExpression, resulting in a total of 905 transcriptomes from 11 datasets. A meta-analytic regularized regression model (“glmnet”, caret package) was trained jointly on all cell types from all studies to use log-normalized gene expression (10,957 genes) to predict age. Performance was evaluated in held out studies (leave study out cross validation) and held out cell types (leave cell type out cross validation, **Fig 3D**). By nature of regularization, this model selects 462 genes out of 10,957 provided genes to predict age robustly across cell types and datasets. Coefficients of genes used in the model are provided in **Supplementary table S4**. The robustness of this cell type-agnostic model was further validated in two datasets (Herb and Zhou) that were held out during training. This model is the one used to evaluate developmental progression in external human neural organoid datasets (Fig 7).

#### Cell type specific models

Initially, we trained cell type-specific models on the transcriptomes of each broad cell type (Excitatory neurons, Inhibitory neurons, Oligodendrocytes, Progenitors, Astrocytes, Vascular, Immune, and Others). These models showed similar performance on their training cell type (MAE in cross-validation) and held-out cell types, indicating limited cell type specificity (Fig 3C). Since their performance was lower than the cell type-agnostic model, they were not used for predictions on external datasets.

### Aggregate co-expression network and module analysis

To characterize gene sets contributing to the cell type agnostic model’s performance, we constructed aggregate co-expression networks for the positive and negative coefficient genes used in the model. First, we constructed a co-expression matrix per dataset using Spearman correlation coefficient between genes calculated from the cell type-level aggregated transcriptomes. The correlation coefficients were then ranked and divided by the maximum rank to obtain a rank-standardized co-expression matrix. Average rank-standardized co-expression across datasets (n =11 used in model training) was computed for each gene-pair to generate the aggregate co-expression network for a set of genes. The aggregate co-expression network was hierarchically clustered using hclust and cut at a specified height using cutree (h = 4.5, or h = 3 for positive and negative gene modules respectively) to generate 9 positive and 11 negative gene modules. The genes corresponding to each module are listed in **Supplementary table S6**.

#### Study-specific age-correlated genes

We identified age-correlated genes within each study by calculating the pearson correlation coefficient and p-value (cor.test) between each gene’s log normalized expression from cell type-aggregated transcriptomes and sample age. P-values were corrected for multiple hypothesis testing using the false discovery rate (FDR) method with p.adjust (method = “fdr”). Statistically significant age-correlated genes (adjusted *P* < 0.05) from each study are listed in **Supplementary table S5.** Enrichment of age-correlated genes from each study in the cell type agnostic model (Fig 4) was calculated using Fisher’s exact test with FDR correction.

### GO term enrichment and characterization

#### Enrichment of GO terms in co-expression modules

GO annotations were obtained from GO.db and org.Hs.eg.db (sourced 2024-01-17). The hypergeometric test (phyper) was used to test the enrichment of specific GO terms in each co-expression module. P-values were corrected for multiple hypothesis testing using the false discovery rate (FDR) method with p.adjust (method = “fdr”). Only GO terms with 10 to 200 genes were evaluated. Results for GO Biological Process terms are shown in Figure 5. Enrichment of GO Biological Process, Cellular Component, and Molecular Function terms in positive and negative co-expression modules are listed in **Supplementary table S7**.

#### Age-prediction performance of specific GO terms

To further explore gene sets driving age-prediction performance across datasets, we trained cell type agnostic and cell type-specific models using regularized regression (caret package, method =glmnet) with genes restricted to specific GO terms. In total, 2213 GO terms with 50-500 genes were tested. As before, cell type-level aggregate transcriptomes were used for training the models. The performances (MAE) of 2213 GO terms for each of the cell type-specific and cell type-agnostic models are shown in **Fig 6** and listed in **Supplementary table S8.** For comparison, models were also trained using the synGO geneset with 1112 genes ^46^. Spearman correlation coefficient of performance was computed between pairs of cell types across all 2213 GO terms to assess cell type-specificity of GO terms in age prediction (Fig 6D).

### Developmental age predictions in external datasets

#### Organoid age predictions

Single cell transcriptomes from human neural organoid datasets (Uzquiano et al, Fiorenzano et al, Revah et al.) were aggregated by author-assigned cell types within each sample using Seurat’s AggregateExpression. The pre-trained cell type agnostic model (Fig 3) predicted ages for each dataset using the predict function from the caret package. Predicted human gestational ages of cell types per organoid sample were compared to organoid *in vitro* culture age (Fig 7A). In the Paulsen et al. dataset, age predictions by the cell type-agnostic model were used to assess differences in the GABAergic lineage between wildtype and ASD-mutant organoids (*ARID1B, SUV420H1, CHD8*). Age differences for all lineages by cell line, gene, and organoid age are shown in **Supplementary** Figs 4-6.

#### Assessing whole-transcriptome similarity with predicted age

Spearman correlation coefficients between cell type-aggregated transcriptomes were computed within each organoid dataset (Uzquiano et al, Fiorenzano et al, Revah et al.), producing a cell-cell similarity matrix based on all genes. To evaluate performance in predicting cell-pair similarity based on age differences, we ranked the correlation coefficients and compared the average rank of pairs with age difference below and above a certain threshold. This results in the Mann-Whitney U-statistic or AUROC, which we computed separately for predicted age and actual organoid age, in cell pairs of the same cell type and different cell types.

#### Age prediction in integrated human neural organoid cell atlas

To support generalization of the cell type-agnostic model to large integrated atlases with varying read depths, we trained a normalization-robust model that uses rank-normalized data instead of log-normalized expression. Here, each gene’s raw count is replaced by its rank relative to other genes within that cell and divided by the maximum rank to obtain rank-normalized expression values within each dataset. The integrated HNOCA h5ad object ^27^ comprising 36 organoid single-cell datasets with 1.77 million cells was downloaded from CellxGene. We aggregated expression at the cell type-level within each dataset and batch, using grouping variables based on ‘publication’, ‘batch’, ‘cell_type_original’, ‘organoid_age_days’,’ state_exact’, and ‘annot_level_1’. This resulted in a total of 9811 transcriptomes, which were rank-normalized and then used for age prediction with the normalization-robust cell type-agnostic model. Predicted ages across all 36 datasets were compared to organoid *in vitro* culture age (**Supplementary** Fig 7).

#### Age predictions in the embryonic mouse brain

The embryonic mouse brain dataset from La Manno et al. with 292,465 cells from 93 samples was downloaded as a loom file and converted to a Seurat object. We aggregated expression at the cell type-level within each sample, resulting in a total of 1177 transcriptomes. We predicted age in the mouse brain using a variant of the cell type-agnostic model trained purely on human brain tissue, but with genes restricted to mouse orthologs present in the La Manno dataset (n = 9460 genes). One to one orthologs between human and mouse were downloaded from http://www.ensembl.org/biomart/martview. Predicted human age was obtained using log-normalized expression of one-to-one mouse orthologs in the embryonic mouse brain dataset and compared to the actual mouse age (embryonic days).

### Compositional vs Cell autonomous predictions

Using predictions from the cell type-agnostic model, we compared the average age of astrocytes, progenitors, and neurons per sample to the actual age of the tissue (embryonic/ *in vitro* days) across mouse brain, Uzquiano et al. organoid, and fetal human brain datasets. Similarly, compositional age for each organoid, human, or mouse brain sample was predicted using the astrocyte-progenitor meta-analytic model and compared to the actual age of the sample. Slopes were computed for each of the curves by fitting a simple linear model. Predicted compositional and cell autonomous ages for each sample were also compared to each other and ANOVA was used to estimate the influence of tissue on developmental dynamics.

### R and R packages

All analysis was carried out in R version 4.4.1. Colors for heatmaps were selected using RColorBrewer and circlize. All plots were made using ggplot2 (v 3.5.1). Heatmaps are plotted using ComplexHeatmap ^72^. All code used to generate figures will be made available at the time of publication. Cell type-agnostic age prediction models are made available at https://github.com/sridevi96/NeuroDevTime.

