## Supplemental figures for "Cell Type-Agnostic Transcriptomic Signatures Enable Uniform Comparisons of Neurodevelopment"

### Supplementary Figures 1-7

Cell type-Agnostic Transcriptomic Signatures Enable Uniform Comparisons of Neurodevelopment

Sridevi Venkatesan, Jonathan M. Werner, Yun Li, Jesse Gillis

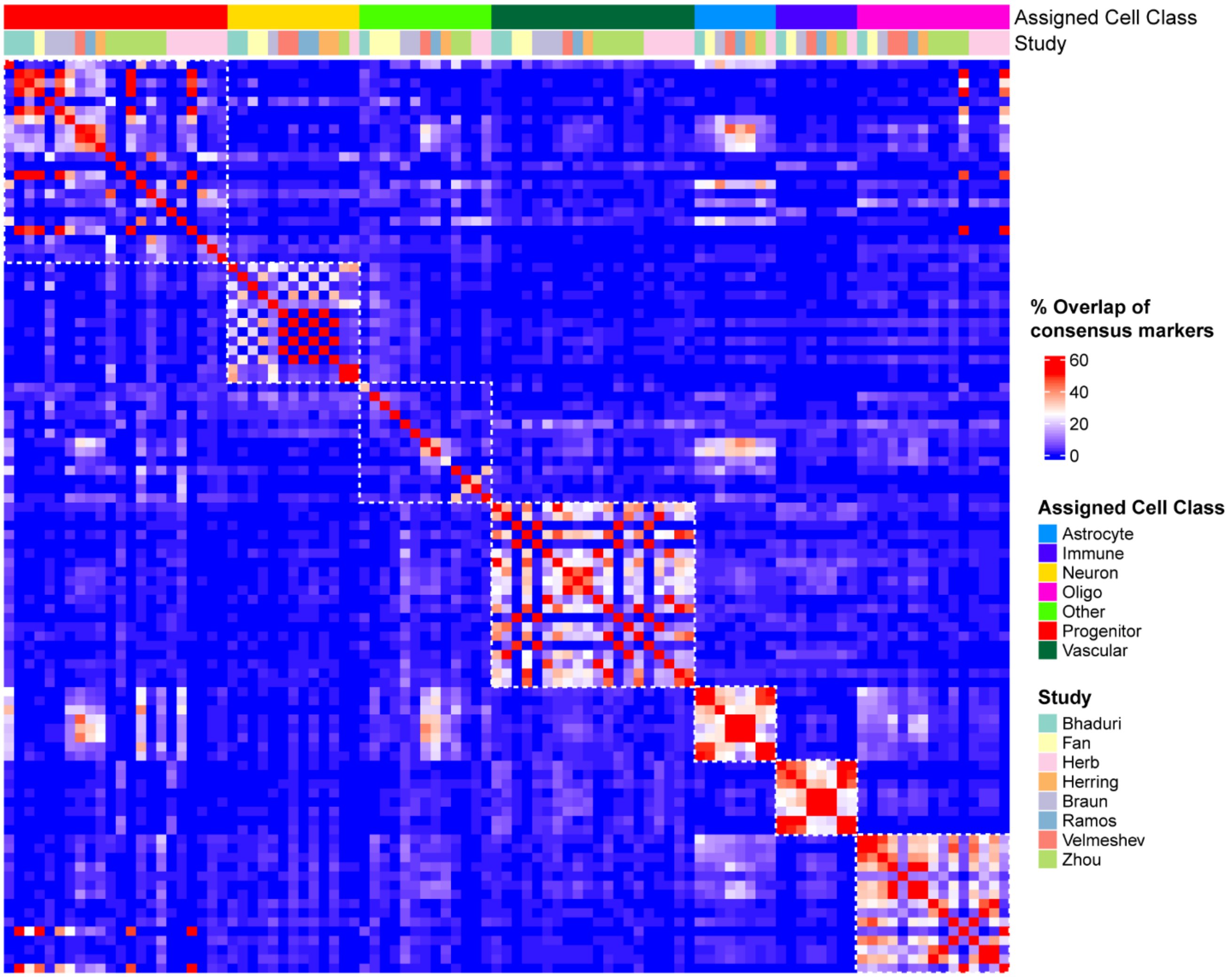


**Supplementary Figure 1: Overlap of consensus marker genes in cell types across human brain datasets**

Author-provided cell type annotations from each study were grouped into 7 broad cell types (Astrocyte, Progenitor, Neuron, Oligodendrocyte, Immune, Vascular, Others; Supplementary table 2). Heatmap shows the overlap of top 100 consensus marker genes for each cell type across datasets. Cell classes show consistent overlaps in consensus markers across datasets, except for “Other” cells which include multiple different cell types.

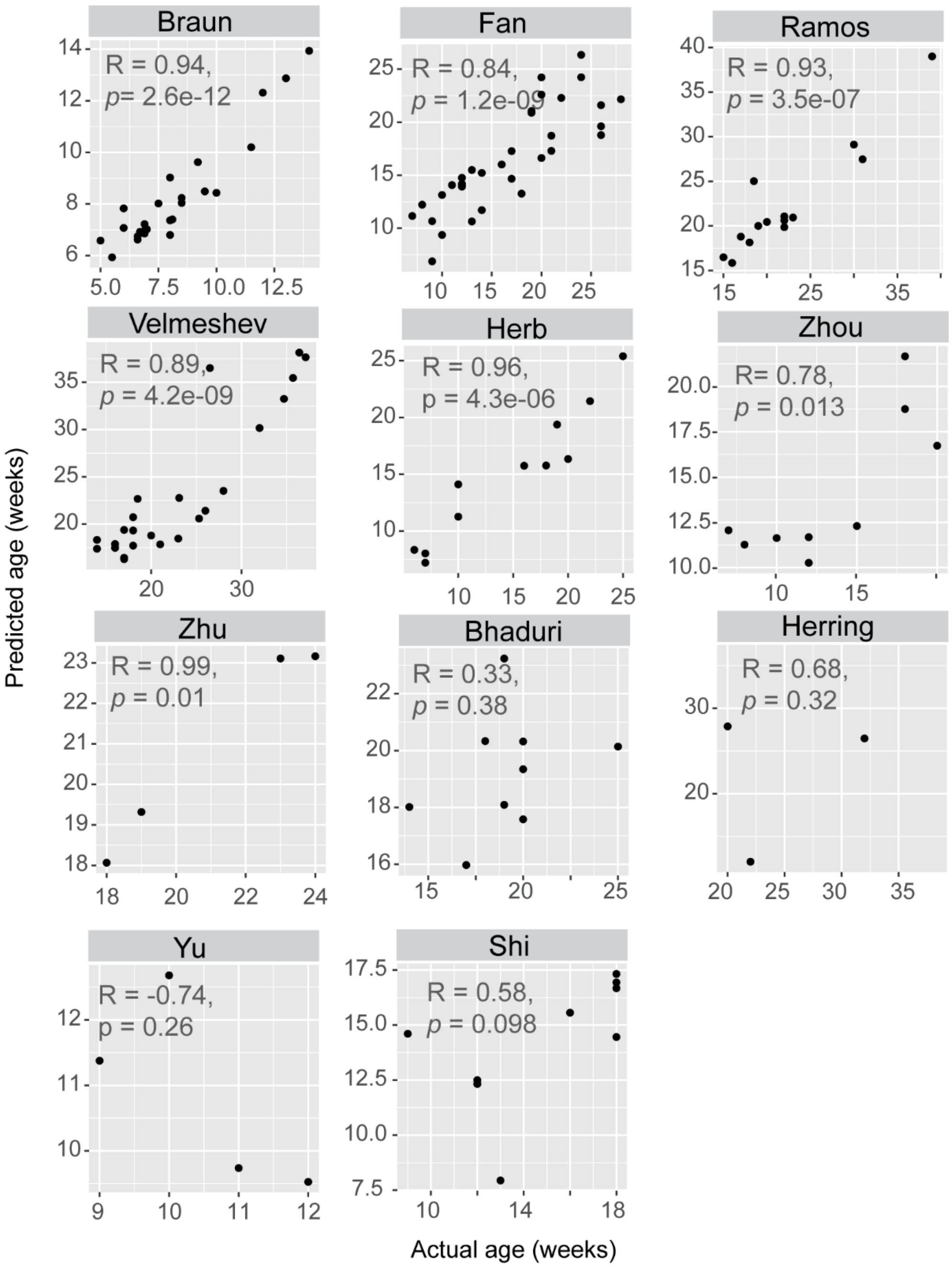


**Supplementary Figure 2: Study specific compositional model performance**

Performance of regularized regression models trained to predict sample age from cell class proportions within each study. Study-specific compositional models accurately predict gestational age in 7 out of 11 studies, with significant correlation between predicted and actual ages.

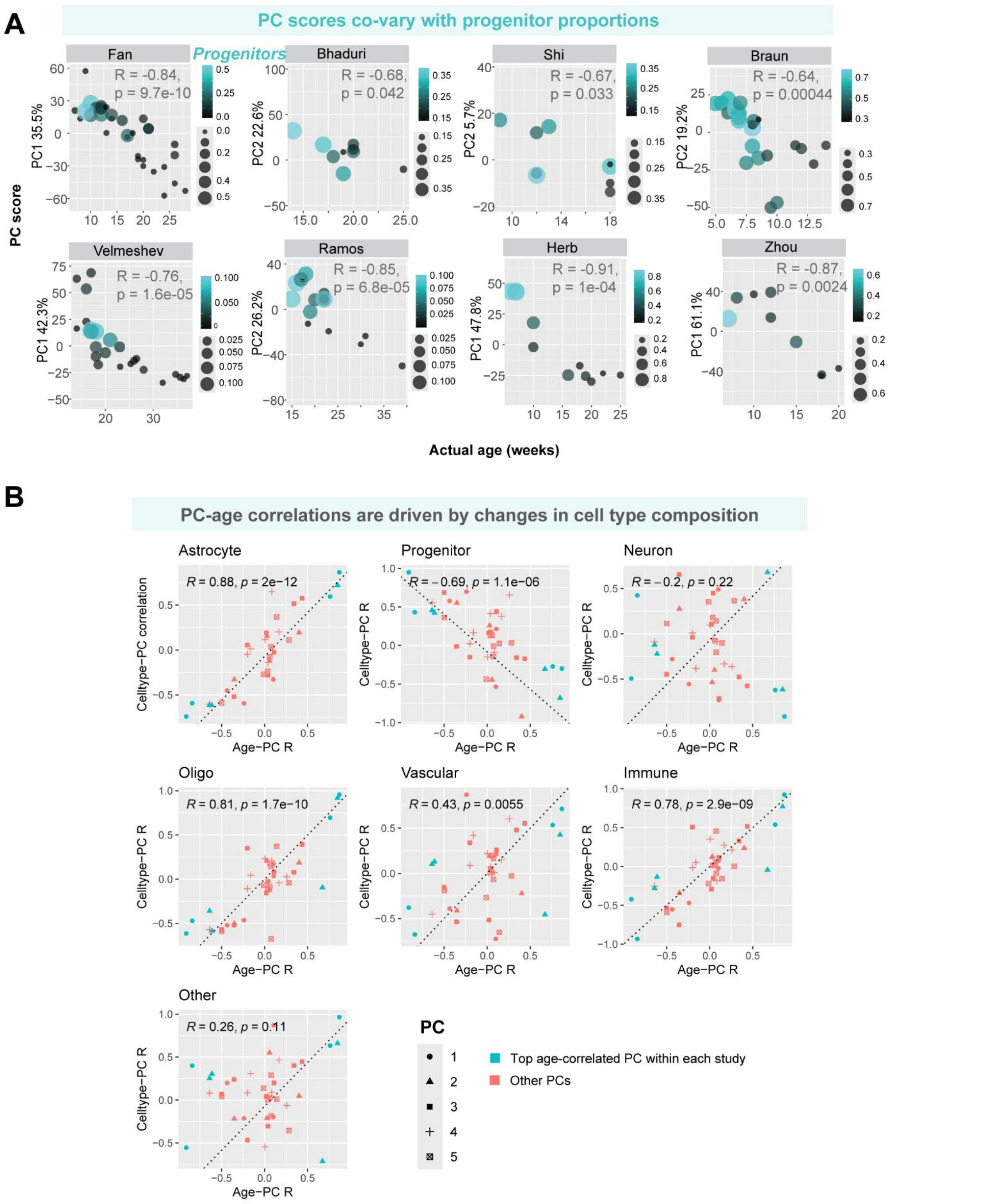


**Supplementary Figure 3: Principal component age predictions are primarily driven by cell type composition.**

**A,** PC1 or PC2 scores are strongly correlated to age within each of the 8 datasets. Size of dots represents the proportion of progenitor cells in the sample. Percentage of variance explained by the PC is shown on the y-axis label. **B,** Correlation of PC with age is plotted against correlation of the PC with proportions for each cell type. Age-PC correlations are perfectly matched by PC-cell type proportion correlations for most cell types across all datasets. Shape of points indicates which principal component, and color represents the PC that was used for age-prediction in panel A.

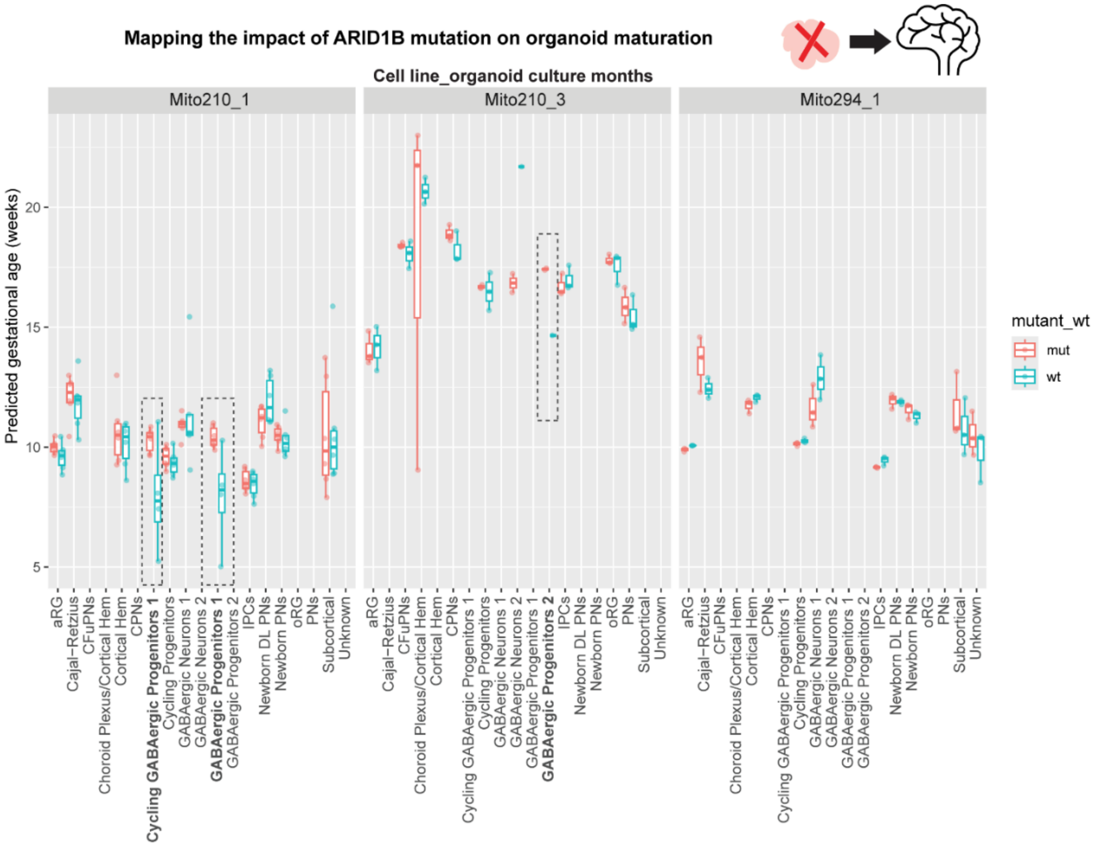
Mapping the impact of ARID1B mutation on organoid maturation
Cell line_organoid culture months
Mito210_1
Mito210_3
Mito294_1
Predicted gestational age (weeks)
5
10
15
20
mutant_wt
mut
wt
aRG
Cajal−Retzius
CFuPNs
Choroid Plexus/Cortical Hem
Cortical Hem
CPNs
Cycling GABAergic Progenitors 1
Cycling Progenitors
GABAergic Neurons 1
GABAergic Neurons 2
GABAergic Progenitors 1
GABAergic Progenitors 2
IPCs
Newborn DL PNs
Newborn PNs
oRG
PNs
Subcortical
Unknown

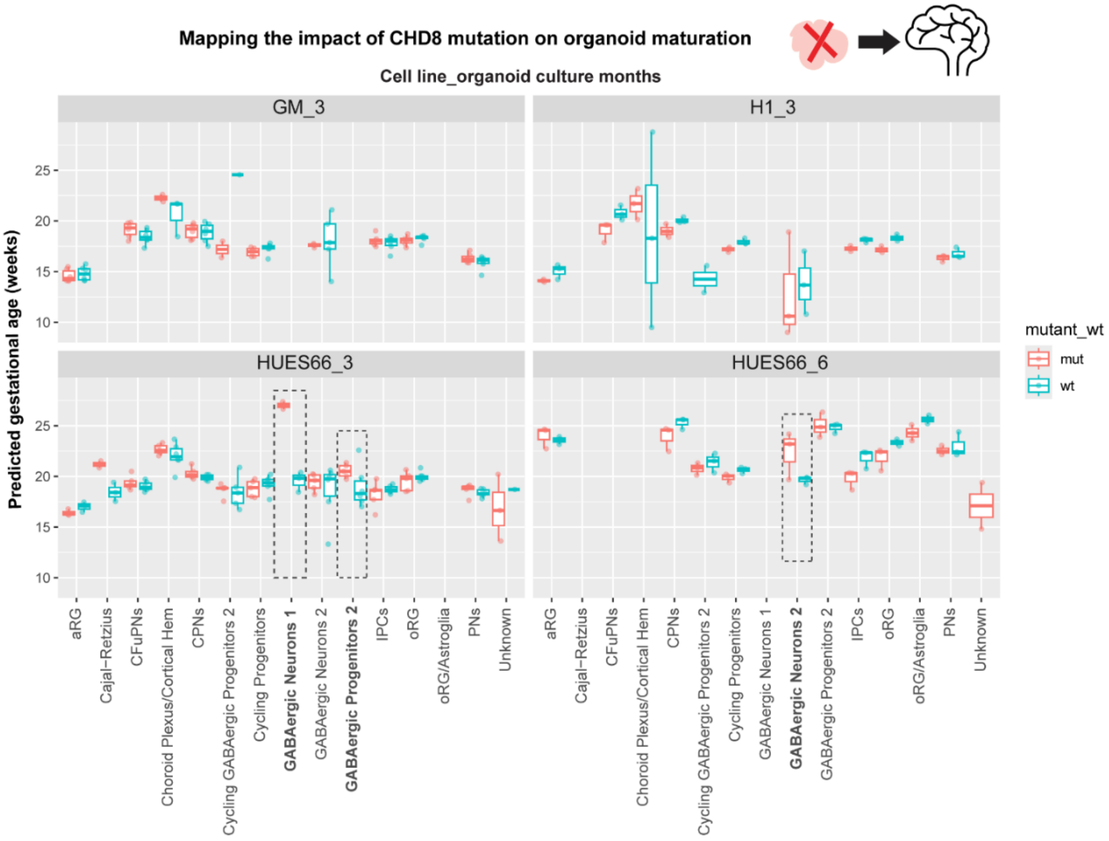
Mapping the impact of CHD8 mutation on organoid maturation
Cell line_organoid culture months
GM_3
H1_3
HUES66_3
HUES66_6
Predicted gestational age (weeks)
10
15
20
25
mutant_wt
mut
wt
aRG
Cajal−Retzius
CFuPNs
Choroid Plexus/Cortical Hem
CPNs
Cycling GABAergic Progenitors 2
Cycling Progenitors
GABAergic Neurons 1
GABAergic Neurons 2
GABAergic Progenitors 2
IPCs
oRG
oRG/Astroglia
PNs
Unknown

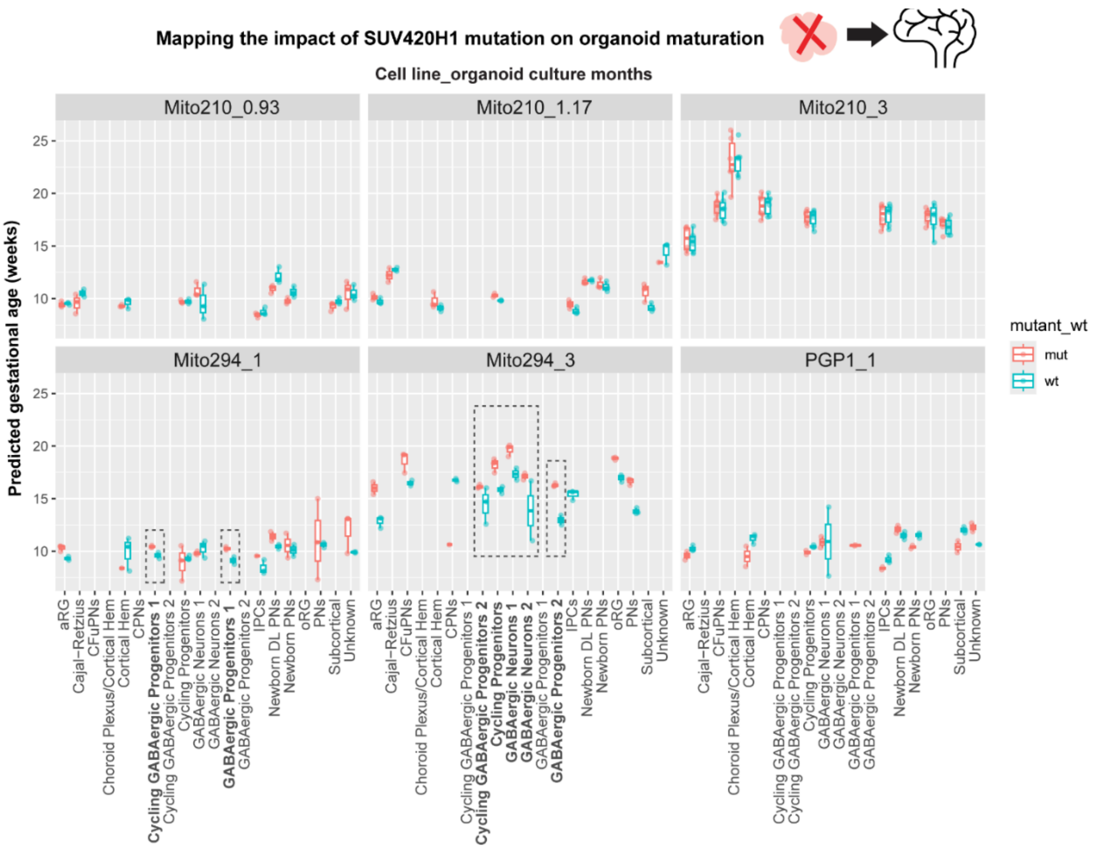


**Supplementary Figures 4-6 Cell autonomous age predictions in ARIDB1, CHD8, and SUV420H1 mutant organoids from Paulsen et al. 2022**

Predicted ages in each cell type from Paulsen et al. 2022 neural organoids separated by gene, cell line, and organoid culture age (in months). Dotted boxes highlight GABAergic cell types where mutant cells show accelerated maturation, as per the original study.

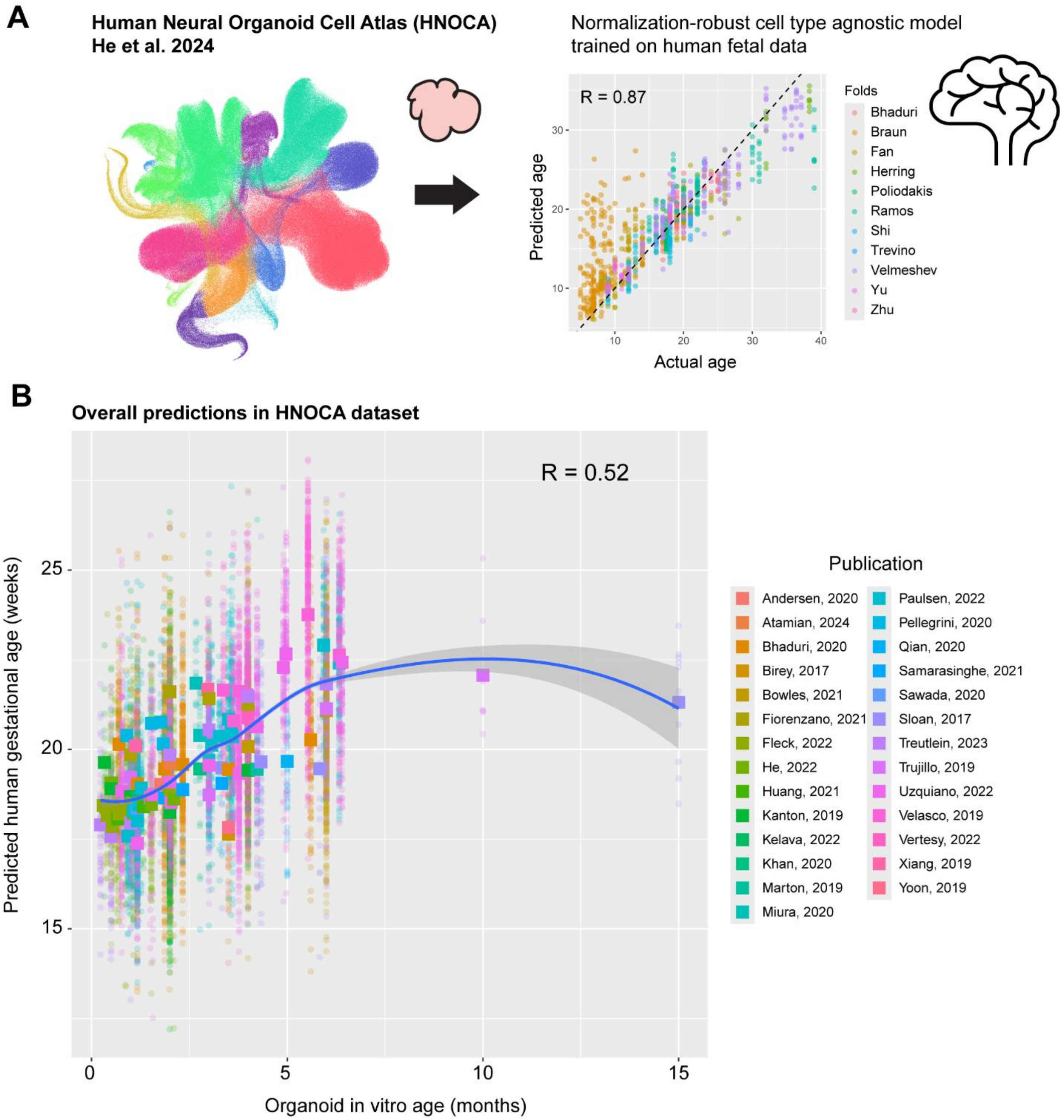


**Supplementary Figure 7: Developmental age predictions in integrated human neural organoid cell atlas (HNOCA)**

**A,** UMAP of HNOCA cells taken from CellxGene browser (left). Right panel depicts cross-validation performance of cell type agnostic model trained on rank-normalized expression values from human fetal brain cell types. **B,** Developmental age predictions for organoid cells from different studies in the integrated HNOCA dataset. Predicted age is overall strongly correlated to organoid culture age *in vitro*. Square points show mean predicted age of all cell types per time point in each study.
